# Finding the known unknowns: minimal machine learning models of resistance identify novel antibiotic resistance discovery opportunities in *Klebsiella pneumoniae*

**DOI:** 10.1101/2025.04.08.647753

**Authors:** Kristina Kordova, Caitlin Collins, Julian Parkhill

## Abstract

Bacterial antimicrobial resistance (AMR) poses a significant public health threat. The advent of global awareness and affordable whole genome sequencing has yielded an ever-growing collection of bacterial genome sequence datasets and corresponding antibiotic resistance metadata. This enables the use of computational techniques, including machine learning (ML), to predict phenotypes and discover novel AMR-associated variants. With the great variety of resistance mechanisms to interrogate and the number of datasets that can be mined, there is a need to identify where novel AMR marker discovery is most necessary. Multiple databases and annotation pipelines exist to identify AMR variants known to be associated with resistance to specific antibiotics or antibiotic classes, however, the completeness of these databases varies and for some antibiotics even the most complete databases remain insufficient for accurate classification. Here, we couple these pipelines with predictive ML models, which we call “minimal models” of resistance. We predict the binary resistance phenotypes of 20 major antimicrobials in the genomically diverse pathogen *Klebsiella pneumoniae*. We present a detailed comparison of the annotation pipelines and drug resistance databases currently available, and we identify their shortcomings in phenotype prediction, highlighting opportunities for novel marker discovery. We further provide a description of a Bacterial and Viral Bioinformatics Resource Center (BV-BRC) database, highlighting the observed AMR mechanism as the key for phenotype prediction in this dataset. This analysis has relevance for all those seeking to use or improve drug resistance databases. It provides a critical review of the differences in annotation tools and databases commonly used in bacterial AMR studies, identifying existing gaps and novel AMR marker discovery niches. It outlines guidance for the establishment of a real standard dataset for the development and benchmarking of ML models of AMR.

## Introduction

As whole-genome sequencing and high-performance computing costs decline, it has become common to utilise increasingly complex *in silico* approaches to predict the antimicrobial resistance (AMR) profile of bacterial isolates. Studies have sought to incorporate genetic variation across the entire genome through the use of known AMR markers, single nucleotide polymorphisms (SNPs), k-mers or unitigs, each providing ever more comprehensive coverage. With the increased use of machine learning (ML), the complexity of prediction algorithms has also soared, building on simpler linear models to develop ensemble and deep learning models, able to disentangle non-linear variant interactions.

As well as high accuracy in predictions, complex models incorporating variation across the bacterial genome have also yielded large numbers of previously uncharacterised variants associated with resistance (Avershina et al. 2021; Liu et al. 2021). The identification of novel variants is believed to be most important in open pangenomes which acquire novel variation more rapidly, such as the genome of *Klebsiella pneumoniae* (Wyres & Holt, 2016). This bacterium has been shown to play a pivotal role in amplifying and shuttling resistance genes across Enterobacteriaceae, hence, discoveries in *K. pneumoniae* could have implications beyond this species (Wyres & Holt, 2018). However, alongside essential discoveries, many of the novel hits can appear spurious due to the rise of high dimensionality and feature correlation challenges as more comprehensive genomic variation is being analysed. Hence, as the possibility of increasing complexity becomes limitless, a question arises of whether increased computing costs and analysis time will always yield biologically meaningful improvements in classification and variant discovery.

We propose that a “minimal model” of resistance can be utilised to begin addressing this enquiry and demonstrate where discovery is most necessary. We define this approach as relying on the known repertoire of AMR genes and mutations previously described as contributors towards resistance to a particular antibiotic or class of antibiotics. In contrast to comprehensive models, what we call “minimal models” are highly computationally efficient, as they take advantage of rapid annotation tools and curated databases (Feldgarden et al. 2019; Jiang et al. 2024). By focusing on well-characterised resistance genes in the diverse *K. pneumoniae* population, coupled with predictive machine learning models, we aim to determine where the minimal model approach significantly underperforms, pointing to the need for the discovery of new AMR mechanisms or variants.

The choices of an annotation tool and reference database are a major determinant of the performance of a minimal model. Multiple databases and computational tools have been published, each curated to enable the annotation of bacterial genomes against known AMR markers. Databases include The Comprehensive Antibiotic Resistance Database (CARD), NCBI’s Reference Gene Catalog, UNIPROT, ARDB, ResFinder, PointFinder, ResFams and others (Alcock et al. 2023; Bortolaila et al, 2020; Gibson et al., 2015; Liu and Pop, 2009; UniProt Consortium et al. 2023; https://www.ncbi.nlm.nih.gov/pathogens/refgene/). While all databases incorporate a list of genes and gene families, only some extend to species-specific point mutations as well, such as PointFinder and the database associated with the annotation tool AMRFinderPlus (Feldgarden et al. 2021). Each database has also been curated with different rules, resulting in differences in antimicrobial resistance gene (ARG) content (Maboni et al. 2022). Some have focused on stringent validation, such as CARD, while others have included an array of variants predicted to have an impact on phenotype with high confidence, such as DeepARG (Arango-Argoty et al, 2018).

Currently, there are more than 18 commonly used species-agnostic open source command line tools designed to rapidly identify the presence of genes and/or mutations (Mendes et al, 2024). The annotation tools such as ABRicate, Resistance Gene Identifier (RGI), AMRFinder and its newer version AMRFinderPlus rely on the previously listed databases to match genomic sequences against the reference and annotate the presence of AMR variants in each sample (Feldgarden et al. 2021; McArthur et al. 2013; https://github.com/tseemann/abricate). Furthermore, species-specific annotation tools have also been developed, which match sequences against the variation reported in the bacteria of interest. These tools have the potential to yield less spurious and more concise gene matching. For example, the tool Kleborate is designed specifically to catalogue the variation in the bacteria *K. pneumoniae* (Lam et al. 2021). Other pipelines include TBprofiler, specialising in annotations of *Mycobacterium tuberculosis* and Mycrobe, focusing on multiple selected species (Verboven et al. 2022; Hunt et al. 2019). The many developed pipelines differ in supported inputs, search algorithms, parameterisation and output formats. The complexity of underlying methodologies also varies significantly, which results in different granularity and quality of annotation. For example, the commonly used Abricate uses the NCBI database by default but only covers a subset of what AMRFinderPlus encompasses, resulting in the inability to detect point mutations, as well as some genes. Hence, an informed choice of annotation tool and database can be difficult and potentially case-specific, inevitably leading to variation in the performance of minimal models (Doyle et al. 2020).

Here, we compare the completeness of gene annotations produced by eight commonly used annotation tools, applied to assembled genomes of *K. pneumoniae*. By doing so, we aim to highlight antibiotics where known mechanisms do not fully account for the observed variation in resistance, and therefore, where ML-enabled marker discovery will be most valuable. We compare pipeline utility in predicting sample phenotype using their defined sets of AMR genes and mutations, which we term “minimal models”. We satisfy minimality by subsetting AMR genes specific to each antimicrobial of interest or its class, as defined by each annotation tool. To generate phenotype predictions from these collections of annotated genes, we used ML models of varying complexity based on the presence/absence of the annotated markers. The performance of each tool is then explained in the context of genes that were overweight by models during prediction. A review of known AMR mechanisms and determinants of resistance to selected antimicrobials is also provided, showing which antibiotics emerge as avenues of interest, either as well-studied or underperforming in classification based on known mechanisms.

## Methods

### Data sourcing and pre-processing

For this analysis, we sought to explore a dataset which is becoming more commonly used as a standard source of bacterial samples - the Bacterial and Viral Bioinformatics Resource Centre (BV-BRC) public database (Olson et al., 2023). The whole genome sequences of 18,645 *K. pneumoniae* samples filtered for good-quality assemblies were obtained. Length of genomes averaged 5.6Mbp and assemblies consisted of up to 1000 contigs. The majority of deposited samples were collected from clinical studies, from both patients and hospital environments, spanning 66 countries. Outlier genomes with more than 250 contigs and lengths of more than 6.4Mbp and less than 4.9Mbp base pairs were excluded from downstream analysis due to low quality and possible contamination. To check for genomes from other species, all sequences were species- and MLST-typed using Kleborate v2.2.0 (Lam et al., 2021). 125 samples were removed, matching to species *K. quasipneumoniae subspecies quasipneumoniae, K. quasipneumoniae subspecies similipneumoniae, K. quasivariicola* and *K. variicola subspecies variicola*.

Antimicrobial resistance data for 76 antibiotics tested by clinical phenotyping was available for 4,976 of the BV-BRC associated genomes, grouping into 15 antibiotic classes. Antibiotics where data was available for less than 1800 samples were excluded from analysis, as the low sample size is expected to yield spurious results. This led to a further reduction of the number of genomes to 3,751. Antibiotic abbreviations are defined in Supplementary Table 1. Although MIC prediction has higher clinical relevance, binary resistance profiles tend to yield higher prediction performance and be more suitable for basic ML architectures (Nguyen et al., 2018). We acknowledge that resistance breakpoint, as reported by the European Committee on Antimicrobial Susceptibility Testing (EUCAST) and Clinical and Laboratory Standards Institute (CLSI) could have changed through the years and re-conversion of the MIC might be beneficial. However, not all studies reported MICs but only binary labels, hence, we utilised the resistance labels as provided by the BV-BRC database to allow for consistency with the standard database. Phenotypes across antibiotics are shown in Figure 1.

**Figure 1.**
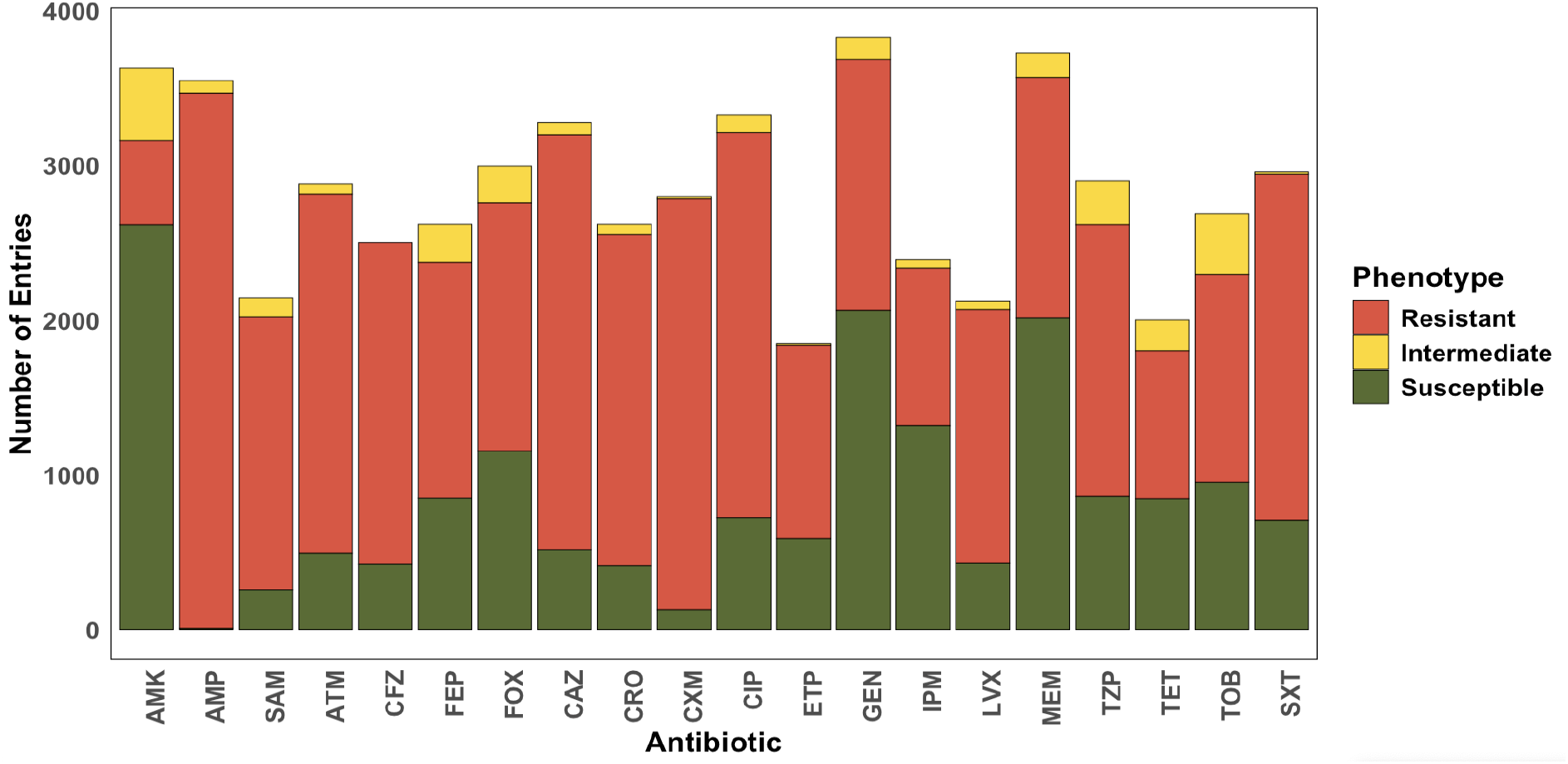
Distribution of phenotypes across antibiotics

### Sample annotation and minimal gene subsets

We reviewed 19 popular annotation tools, discussed by previous studies (Supplementary Table 2), of which eight were applicable to the analysis of assembled *K. pneumoniae* genomes (Table 1) (Mendes et al., 2024). After review, the available samples were annotated using Kleborate, ResFinder, AMRFinderPlus and DeepARG against their default database settings, RGI, SraX and Abricate against the CARD database, and StarAMR against ResFinder. Virulence gene hits were excluded from these annotations for the purpose of building a minimal model of antimicrobial resistance. The tools annotated samples in gene-to-antibiotic and gene-to-class relationships. To compare the performance of minimal models predicting resistance to antibiotic classes with resistance to individual antibiotics, we further acquired a list of genes corresponding to each of the 19 antibiotics of interest from the stringent CARD ontology database. The database encompasses gene-to-antibiotic and mutation-to-antibiotic relationships which have shown experimental increase in MIC. Subsets of genes-to-class and genes-to-antibiotic were used for downstream ML model fitting. In cases where susceptibility testing was carried out for combination therapies, the genes for both classes were included in the minimal subset (e.g. the gene set for amoxicillin-tetracycline was a combination of the sets for amoxicillin and tetracycline). Where genes were annotated as multi-drug, they were included in all subsets, as they marked efflux pups and porins relevant across antibiotics. These were not subset further as their specific function in relation to each class or antibiotic is difficult to disentangle or is not fully known. No additional feature engineering was carried out and annotations were used as they were produced.

**Table 1:**
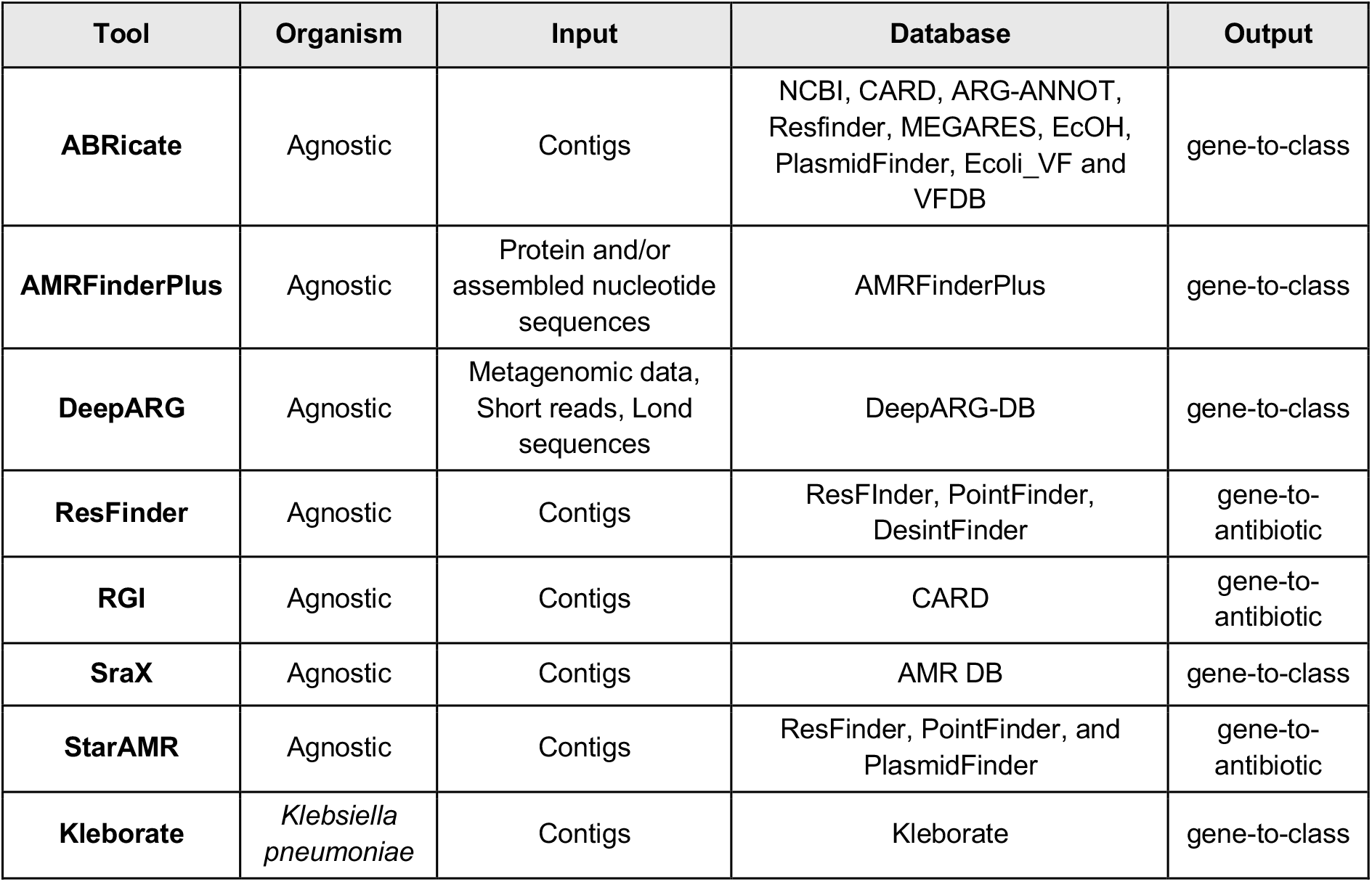
Eight AMR gene annotation tools which were compared in this analysis, describing their target species, input, supported databases and output format of the relationship between the gene and the associated AMR by either antibiotic or antibiotic class..

### Machine learning models

The performance of two types of predictive models was compared when using the provided minimal marker subsets as features, namely logistic regression with L1 and L2 regularisation (Elastic Net) and the Extreme Gradient Boosted ensemble model (XGBoost). These were chosen due to their interpretability and scalability as well as usually high accuracy. Their specific characteristics and parameters are described in the following section. The dataset was split into 70% training and 30% testing, using a stratified fivefold cross-validation without hyperparameter tuning. The performance of the models was scored by observing the sensitivity, specificity and area under the curve (AUC). Due to imbalances in the relative number of resistant and sensitive isolates across antibiotics, we find that AUC appeared as a more informative reporter of the model fit than the more commonly used balanced accuracy. The target labels were binarised so ‘resistant’ also encompassed ‘intermediate’ samples, as reported by the BV-BRC database.

To explain differences in model performance, we scored the names and number of genes used for prediction by each annotation tool. Where the ARG determinants of classification performance were not immediately obvious, we used Shapley values to explain the XGBoost model and logistic regression coefficients to identify features which received high-importance scores in classifications. Shapley values were extracted using the SHAP python package (Lundberg and Lee, 2017). They represent the contribution of each feature in all possible feature combinations in our ensemble XGBoost models. Important feature sets were then compared between tools which show high and low performance for the antibiotic of interest.

### Elastic Net

Elastic Net is a penalised linear regression model that includes both the L1 (LASSO) and L2 (Ridge) penalties in the loss function during training. The L1 penalty shrinks the coefficients towards zero, which removes many predictors from the model. The L2 penalty instead minimises the Euclidean norm of the coefficient vector but typically produces models that use all the predictors. The model assumes a linear relationship between input variables and outputs, hence omitting the possibility of potential inter-dependencies between multiple variants and the phenotype. Elastic Net is believed to manage dependent variables, expected due to the strong linkage disequilibrium in bacterial populations (Biffignandi et al. 2024). The model was tested using α = 0.01 and a switching parameter of 0.5, indicating the model performs as a Ridge selection and as LASSO. Weight balancing between resistant and susceptible classes was not included in the final report. The model was run using the python package sklearn v4.1.3.

### XGBoost

XGBoost is a tree ensemble method that provides the advantage of decorrelating sets of trees through feature and sample subsampling. Studies have reported XGBoost to manage the multicollinearity of small feature sets (Biffignandi et al. 2024). It allows for feature interaction, as features which form the follow-on nodes on each decision tree depend on the features in the previous nodes. Hence this approach advances the Elastic Net by allowing for an extra level of statistical and potentially biological complexity. The XGBoost model was trained using 30 trees with a depth of 3 to reduce the possible overfitting of the small minimal models. Weight balancing between target classes was not included to yield comparable results to the Elastic Net model. The Random Forest model was run using the python xgboost package v2.1.1.

### RGI

The RGI pipeline is versatile in the supported input formats, covering genomes, genome assemblies, metagenomic contigs, k-mers, reads or proteomes (Alcock et al., 2023). It carries out gene prediction using Prodigal, homolog detection using DIAMOND, and Strict significance based on CARD curated bit score cut-offs (Altschul et al. 1990; Hyat et al., 2010). The tool outputs annotations in a gene-to-antibiotic relationship. We used RGI v.6.0.3.

### ResFinder

The ResFinder pipeline utilises BLAST and offers a choice of annotation databases, namely ResFinder, PoinFinder and DesinFinder (Bortolaia et al. 2020). The ResFinder database curates horizontally acquired ARGs and outputs annotations in a gene-to-antibiotic relationship. On the other hand, PoinFinder annotates putative point mutations in ARGs in a gene-to-class manner. Finally, DesinFinder curates resistance markers associated with disinfectants, which were not included in this analysis. The ResFinder tool also predicts the phenotype of samples for specific antibiotics, however, this functionality was used in this analysis to ensure concordance across predictions made with other tools. We utilised ResFinder v.4.1.5 which was last updated only in 2021 (Bortolaila et al. 2020).

### AMRFinderPlus

AMRFinderPlus is the updated version of AMRFinder, released by NCBI (Feldgarden et al. 2021). The pipeline has the advantage of being able to detect the presence of both genes and point mutations against its own curated database. Sequences are matched using BLAST and Hidden Markov Models. AMRFinderPlus v.3.12.8 was used.

### DeepArg

Based on a pre-trained multiclass deep learning model, DeepArg is able to detect AMR genes from both short and long sequences (Arango-Argoty et al, 2018). The tool collates and curates the databases ARDB, CARD and UNIPROT, significantly increasing the diversity of available known AMR sequences. The prediction is carried out by generating a dissimilarity matrix between sequences from predicted genes and known ARGs using DIAMOND, instead of BLAST, which reports higher speed. The resulting output is genes associated with antibiotic classes. The classification is reported to differ in fidelity between classes, with major antibiotic classes being well classified and classes with fewer known genes and multi-drug resistance gene classes performing poorly. Here, genomes were annotated using DeepArg v.1.0.2. against the corresponding database for this software version. Annotations with a probability greater than 80% were included in the analysis. Genes annotated as ‘multi-drug’ were included in all subsets.

### Abricate

The Abricate tool is designed to detect ARGs from assembled contigs by aligning genomes to reference databases using BLAST (https://github.com/tseemann/abricate). The user can select the reference database from an extensive list, including NCBI, CARD, ARG-ANNOT, and others. Hence, an informed user choice could be necessary to select the relevant database. Furthermore, the tool encompasses only gene presence but not point mutations. Abricate v1.0.0. was used to annotate all available genomes against the CARD database.

### Kleborate

Kleborate uses assemblies and outputs annotations of resistance genes by drug class, alongside other quality control metrics (Lam et al., 2021). Nucleotide BLAST is used against the CARD database, as well as other genes and alleles which have been clinically demonstrated to impact MIC profiles in the *K. pneumoniae* species complex. As well as genes with high identity match, the tool also annotates truncated and spurious hits which were included as features if associated with the antibiotic class of interest. This yields a high number of annotations, as we conserve genes with lower identity matches as separate features from genes with high identity matches and rely on the downstream models to assign the appropriate informativeness. We used Kleborate v.2.2.0 rather than the newer v.3.0.0 due to software compatibility.

### sraX

The sraX tool utilises a locally compiled AMRDB connecting to CARD as the default database (Panunzi, 2020). A homolog search is performed by alignments to the database, facilitated using DIAMOND. The tool annotates genome assemblies and provides a gene-to-class level annotation. The sraX v.1.5 was used and annotation identities and sequence matches were set to 95% and 90% respectively. Genes annotated as ‘multi-drug’ were included in all subsets.

### StarAMR

Staramr was created to query the ResFinder, PointFinder and PlasmidFinder databases using a BLAST-based approach to scan bacterial genome contigs for ARGs and mutations (Bharat et al., 2022). It compiles a summary report of genetic mechanisms and a predicted antibiogram based on a gene-to-antibiotic key. We used the only release of Staramr – v.0.10.0, and the corresponding database updates as of January 2023.

### Not included annotation tools

Reviews and consortiums, such as the Public Health Alliance for Genomic Epidemiology (PHA4GE) have additionally noted a few more annotation tools which were not used for the construction of our minimal models. The popular Ariba, AMR++ and SRST2 pipelines were not used as they use raw reads, instead of genome assemblies (Bonin et al. 2023; Hunt et al. 2017; Inouye et al. 2014). GROOT and KmerResistance were excluded as they use metagenomic data or poorly sequenced genomes respectively (Clausen et al. 2016; Rowe and Winn, 2018). Similarly, fARGene utilises fragmented sequences and uses pre-trained AMR class predictive models to predict full-length sequences, however, models need to be run individually, which was not feasible for our analysis (Berlund et al. 2019). Mycrobe was not included as it does not support models of resistance for *K. pneumoniae* (https://github.com/mykrobe-tools/mykrobe). c-STAR and ResFams were not included due to installation challenges (Bharat et al 2020; Gibson et al., 2014).

## Results

The AMR-associated genes and variants identified by the eight commonly-used annotation pipelines listed in Table 1 were used as features in ML models to predict the drug resistance phenotype of each sample. The aim of this analysis was to determine where these minimal models of drug resistance, containing antibiotic- or antibiotic class-specific AMR markers identified by each tool, fail to accurately predict drug resistance. This comparison serves two purposes. First, in Figure 2, we show which antibiotics remain poorly predicted across all AMR databases and annotation approaches, indicating the need for further study. Second, in Table 2, we reveal that substantial variations in predictive performance exist within as well as across antibiotics, depending on which AMR database and annotation tool is used to define the set of known AMR markers in the minimal model.

**Figure 2.**
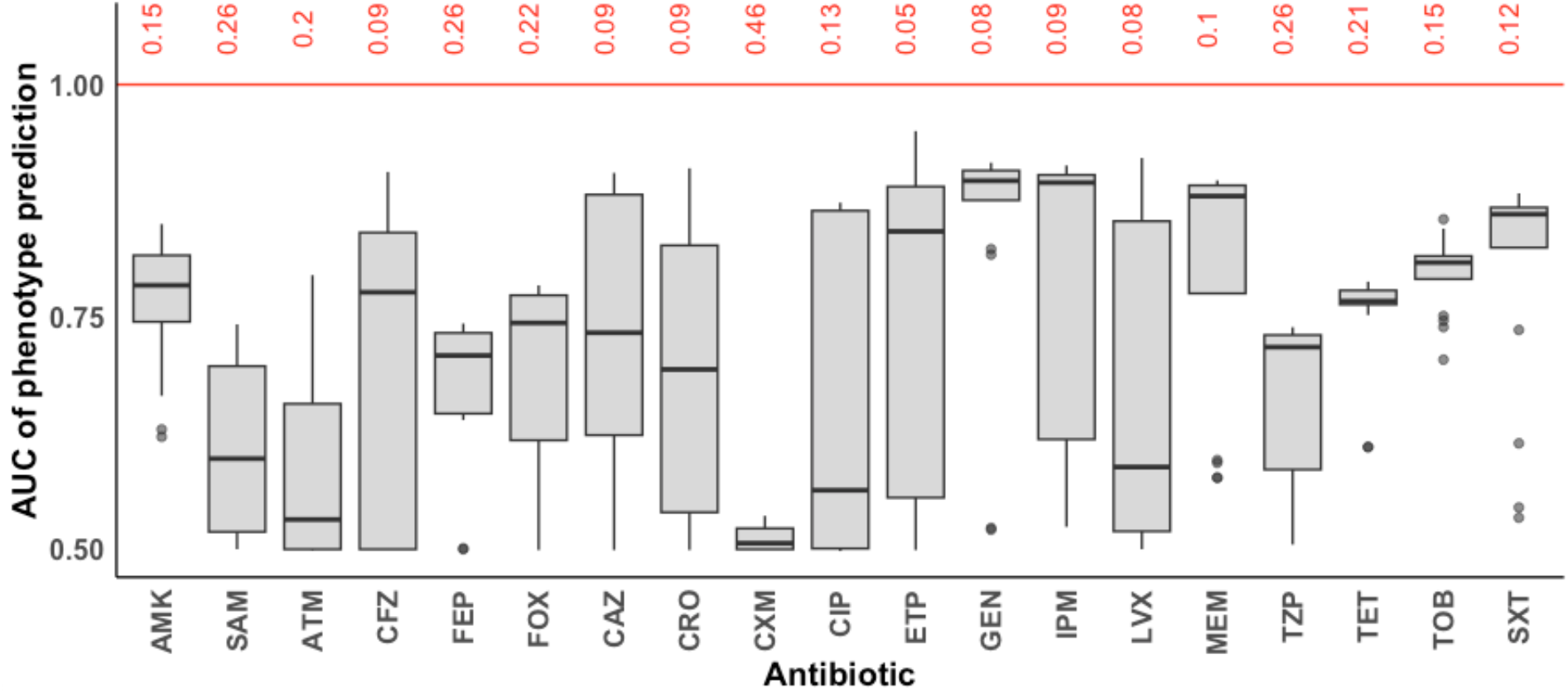
Summary of the minimal model performance across antibiotics showing average performance across annotation databases. The difference between the highest AUC value achieved and one (complete predictive success) is indicated in red for each antibiotic. Outlier predictions AUC value of 0 are excluded as they result from having no annotated features for the respective antibiotic.

**Table 2:**
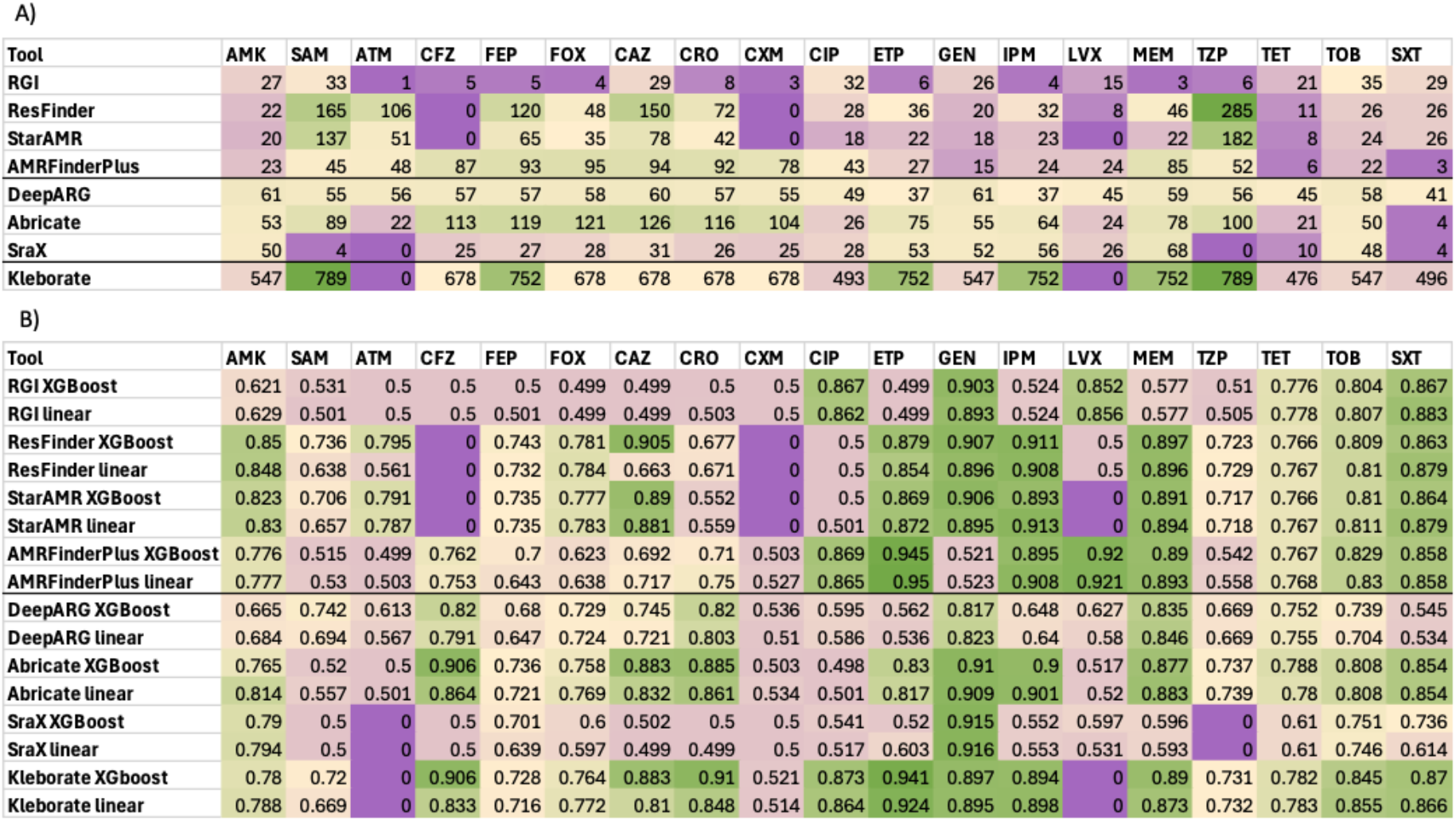
Summary of the annotation and prediction results across antibiotics and annotation tools. A) Number of AMR-associated variants identified by each tool for each antibiotic or antibiotic class and B) AUC across all annotation tools differed significantly. Results are coloured from dark purple (low values) to green (highest values). Bold lines divide tools carrying out gene-to-antibiotic prediction from gene-to-class prediction. Gene number for Kleborate is lined out and coloured separately due to the stark difference in gene number.

| Tool | AMK | SAM | ATM | CFZ | FEP | FOX | CAZ | CRO | CXM | CIP | ETP | GEN | IPM | LVX | MEM | TZP | TET | TOB | SXT |
| --- | --- | --- | --- | --- | --- | --- | --- | --- | --- | --- | --- | --- | --- | --- | --- | --- | --- | --- | --- |
| RGI | 27 | 33 | 1 | 5 | 5 | 4 | 29 | 8 | 3 | 32 | 6 | 26 | 4 | 15 | 3 | 6 | 21 | 35 | 29 |
| ResFinder | 22 | 165 | 106 | 0 | 120 | 48 | 150 | 72 | 0 | 28 | 36 | 20 | 32 | 8 | 46 | 285 | 11 | 26 | 26 |
| StarAMR | 20 | 137 | 51 | 0 | 65 | 35 | 78 | 42 | 0 | 18 | 22 | 18 | 23 | 0 | 22 | 182 | 8 | 24 | 26 |
| AMRFinderPlus | 23 | 45 | 48 | 87 | 93 | 95 | 94 | 92 | 78 | 43 | 27 | 15 | 24 | 24 | 85 | 52 | 6 | 22 | 3 |
| DeepARG | 61 | 55 | 56 | 57 | 57 | 58 | 60 | 57 | 55 | 49 | 37 | 61 | 37 | 45 | 59 | 56 | 45 | 58 | 41 |
| Abricate | 53 | 89 | 22 | 113 | 119 | 121 | 126 | 116 | 104 | 26 | 75 | 55 | 64 | 24 | 78 | 100 | 21 | 50 | 4 |
| SraX | 50 | 4 | 0 | 25 | 27 | 28 | 31 | 26 | 25 | 28 | 53 | 52 | 56 | 26 | 68 | 0 | 10 | 48 | 4 |
| Kleborate | 547 | 789 | 0 | 678 | 752 | 678 | 678 | 678 | 678 | 493 | 752 | 547 | 752 | 0 | 752 | 789 | 476 | 547 | 496 |

Table 2: Summary of the annotation and prediction results across antibiotics and annotation tools.
| Tool | AMK | SAM | ATM | CFZ | FEP | FOX | CAZ | CRO | CXM | CIP | ETP | GEN | IPM | LVX | MEM | TZP | TET | TOB | SXT |
| --- | --- | --- | --- | --- | --- | --- | --- | --- | --- | --- | --- | --- | --- | --- | --- | --- | --- | --- | --- |
| RGI XGBoost | 0.621 | 0.531 | 0.5 | 0.5 | 0.5 | 0.499 | 0.499 | 0.5 | 0.5 | 0.867 | 0.499 | 0.903 | 0.524 | 0.852 | 0.577 | 0.51 | 0.776 | 0.804 | 0.867 |
| RGI linear | 0.629 | 0.501 | 0.5 | 0.5 | 0.501 | 0.499 | 0.499 | 0.503 | 0.5 | 0.862 | 0.499 | 0.893 | 0.524 | 0.856 | 0.577 | 0.505 | 0.778 | 0.807 | 0.883 |
| ResFinder XGBoost | 0.85 | 0.736 | 0.795 | 0 | 0.743 | 0.781 | 0.905 | 0.677 | 0 | 0.5 | 0.879 | 0.907 | 0.911 | 0.5 | 0.897 | 0.723 | 0.766 | 0.809 | 0.863 |
| ResFinder linear | 0.848 | 0.638 | 0.561 | 0 | 0.732 | 0.784 | 0.663 | 0.671 | 0 | 0.5 | 0.854 | 0.896 | 0.908 | 0.5 | 0.896 | 0.729 | 0.767 | 0.81 | 0.879 |
| StarAMR XGBoost | 0.823 | 0.706 | 0.791 | 0 | 0.735 | 0.777 | 0.89 | 0.552 | 0 | 0.5 | 0.869 | 0.906 | 0.893 | 0 | 0.891 | 0.717 | 0.766 | 0.81 | 0.864 |
| StarAMR linear | 0.83 | 0.657 | 0.787 | 0 | 0.735 | 0.783 | 0.881 | 0.559 | 0 | 0.501 | 0.872 | 0.895 | 0.913 | 0 | 0.894 | 0.718 | 0.767 | 0.811 | 0.879 |
| AMRFinderPlus XGBoost | 0.776 | 0.515 | 0.499 | 0.762 | 0.7 | 0.623 | 0.692 | 0.71 | 0.503 | 0.869 | 0.945 | 0.521 | 0.895 | 0.92 | 0.889 | 0.542 | 0.767 | 0.829 | 0.858 |
| AMRFinderPlus linear | 0.777 | 0.53 | 0.503 | 0.753 | 0.643 | 0.638 | 0.717 | 0.75 | 0.527 | 0.865 | 0.95 | 0.523 | 0.908 | 0.921 | 0.893 | 0.558 | 0.768 | 0.83 | 0.858 |
| DeepARG XGBoost | 0.665 | 0.742 | 0.613 | 0.82 | 0.68 | 0.729 | 0.745 | 0.82 | 0.536 | 0.595 | 0.562 | 0.817 | 0.648 | 0.627 | 0.835 | 0.669 | 0.752 | 0.739 | 0.545 |
| DeepARG linear | 0.684 | 0.694 | 0.567 | 0.791 | 0.647 | 0.724 | 0.721 | 0.803 | 0.51 | 0.586 | 0.536 | 0.823 | 0.64 | 0.58 | 0.846 | 0.669 | 0.755 | 0.704 | 0.534 |
| Abricate XGBoost | 0.765 | 0.52 | 0.5 | 0.906 | 0.736 | 0.758 | 0.883 | 0.885 | 0.503 | 0.498 | 0.83 | 0.91 | 0.9 | 0.517 | 0.877 | 0.737 | 0.788 | 0.808 | 0.854 |
| Abricate linear | 0.814 | 0.557 | 0.501 | 0.864 | 0.721 | 0.769 | 0.832 | 0.861 | 0.534 | 0.501 | 0.817 | 0.909 | 0.901 | 0.52 | 0.883 | 0.739 | 0.78 | 0.808 | 0.854 |
| SraX XGBoost | 0.79 | 0.5 | 0 | 0.5 | 0.701 | 0.6 | 0.502 | 0.5 | 0.5 | 0.541 | 0.52 | 0.915 | 0.552 | 0.597 | 0.596 | 0 | 0.61 | 0.751 | 0.736 |
| SraX linear | 0.794 | 0.5 | 0 | 0.5 | 0.639 | 0.597 | 0.499 | 0.499 | 0.5 | 0.517 | 0.603 | 0.916 | 0.553 | 0.531 | 0.593 | 0 | 0.61 | 0.746 | 0.614 |
| Kleborate XGBoost | 0.78 | 0.72 | 0 | 0.906 | 0.728 | 0.764 | 0.883 | 0.91 | 0.521 | 0.873 | 0.941 | 0.897 | 0.894 | 0 | 0.89 | 0.731 | 0.782 | 0.845 | 0.87 |
| Kleborate linear | 0.788 | 0.669 | 0 | 0.833 | 0.716 | 0.772 | 0.81 | 0.848 | 0.514 | 0.864 | 0.924 | 0.895 | 0.898 | 0 | 0.873 | 0.732 | 0.783 | 0.855 | 0.866 |

The ability to predict AMR status from known AMR markers varied considerably across the 19 antibiotics examined. Figure 2 displays the predictive performance of the minimal model for each antibiotic, across all eight annotation tools. Low AUC values indicate where predictions are most discordant from the observed binary resistance phenotype for a particular antibiotic. While AUC values over 0.9 are attained for seven of the 19 antibiotics, none of the 19 antibiotics achieves an average AUC above this level. Resistance to gentamicin (GEN), trimethoprim/sulfamethoxazole (SXT), tetracycline (TET) and tobramycin (TOB) were on average well predicted by most tools, while resistance to cefuroxime (CXM), ampicillin/sulbactam (SAM) and piperacillin/tazobactam (TZP) was most often predicted poorly. While performance appear to vary greatly, AUC values for cefuroxime (CXM), gentamicin (GEN), trimethoprim/sulfamethoxazole (SXT), gentamicin (TET) and tobramycin (TOB) appeared most consistent across tools.

The number of annotated AMR genes and variants identified differed significantly between annotation tools and across antibiotics (Table 2a). RGI was shown to be the most conservative approach across all antimicrobials, identifying the fewest AMR markers across most antibiotics, likely due to the underlying annotation methodology and the gene-to-antibiotic relationship of variants. On the other hand, ResFinder annotated more hits across antibiotics, but failed to identify ARGs associated with resistance to cefazolin (CFZ) and cefuroxime (CXM). AMRFinderPlus, DeepARG and Abricate found a higher number of annotated genes across most antibiotics. Number of genes identified by DeepARG was fairly stable as genes annotated as multi-class were included in all antibiotic-specific subsets. Abricate failed to identify genes for aztreonam (ATM), as the monobactam class was not present in this database, and piperacillin/tazobactam (TZP) as no penicillin or beta-lactam AMR genes were present in the subset of samples. Kleborate typically discovered the greatest number of ARGs, as the tool annotates truncated proteins and genes with different percentage identities, which we kept as distinct features. No associated genes were identified for ampicillin/sulbactam (SAM) and levofloxacin (LVX). Major differences were observed in the phenotype prediction performances of the eight tools by the two ML models (Table 2b, Supplementary Table 3). While in most cases the higher number of predictive AMR markers resulted in higher performance, this was not always the case. The gene-to-antibiotic tools showed distinct performances, with the RGI showing lower AUC than ResFinder for most antibiotics. This is in line with the fewer ARGs annotated by RGI, which limits the prediction potential, while ResFinder relies on a much wider database. All annotated variants per tool and antibiotic are displayed in Supplementary Table 4, as well as the Shapley values demonstrating variants with higher weight in each prediction task.

Significant prediction performance differences were observed for the antibiotics ciprofloxacin (CIP) and levofloxacin (LEV). AMRFinderPlus and Kleborate results contained more granular characterisation of the gyrase genes *gyrA* and *gyrB*, including putative mutations, which likely led to good classification performance. RGI, on the other hand, contained multiple gyrase genes but did not characterise specific mutations. Gyrases were absent in the ResFinder, DeepArg and Abricate annotations, which likely led to the inability to predict the fluoroquinolone resistance profile using these tools. Instead, these tools annotated multiple *qnrb and qnra* genes. The same pattern is observed for levofloxacin (LEV) which also falls into the same antibiotics class. The mechanisms associated with these genes are reviewed in the following section.

Resistance to sulfamethoxazole/trimethoprim (SXT) was well predicted by AMRFinderPlus and Abricate by annotating only three genes, namely *sul1, sul2* and *sul3*. These genes were also annotated by all other tools as well. Interestingly, while these genes were annotated by DeepARG, they were often identified in incorrect samples, which likely led to the underperformance of this tool. *dfrA* gene variants were also given high importance whenever detected by annotation tools, in line with the nature of resistance to combination therapy. Gentamicin (GEN) resistance was poorly predicted by AMRFinderPlus, which was missing annotations for *aac(3)* genes, annotated by all other tools, as well as *armA*. These genes are major determinants of resistance to aminoglycosides and have a high Shapley value for the classification of gentamicin by most tools. The genes were also missing in the amikacin AMRFinderPlus annotation, however, they did not receive a high Shapley value in predictions by other tools. Instead, variants of the genes *aac(6)* and *aph(3’) -VI* were shown to have high importance, which were annotated by all tools.

ML models for aztreonam (ATM), cefepime (FEP), cefoxitin (CFX), piperacillin/tazobactam (TZP), tetracycline (TET) and tobramycin (TOB) performed mostly universally across tools, despite the large differences in gene numbers. In some cases, such as ampicillin/sulbactam (SAM), cefuroxime (CRO) and ceftazidime (CAZ), the XGBoost model overperformed the Elastic Net predictions. This could be interpreted as arising from the higher model complexity or due to the need to incorporate variant interaction as a strong determinant of resistance. However, as the annotations do not provide a granular and comprehensive genome coverage, the role of interaction effects in these antibiotics needs to be studied further.

Resistance to meropenem (MEM) was poorly classified based on the annotation from RGI, likely due to many missing genes, as this tool detected only three AMR genes for this antibiotic. The underperformance in imipenem (IMP) classification by RGI and DeepARG was likely due to the missing annotation of *blaKPC* and *ompK*, which were present in all other annotations and marked as important by high Shapley values.

## Discussion

In this analysis, we demonstrate that annotation tools show stark differences in their annotation patterns and breath, which further translates to a varying and often low utility for ML-enabled phenotype prediction. This is in line with expectations and confirms previous findings of multiple studies, which demonstrate that the use of known ARGs is insufficient to provide clinical-grade testing, where a maximum error rate of 3% is permitted (Doyle et al., 2020; Maboni et al., 2022; Mahfouz et al. 2020). We further demonstrate that no annotation tool is consistently better than the others across all antibiotics, with pipelines such as AMRFinderPlus overperforming in predictions of some antibiotics while underperforming in others. However, annotations with Kleborate showed the most consistent usefulness in ML classifications. This finding is also in line with expectations, as Kleborate is specifically curated to include AMR mechanisms in the bacterium of interest *K. pneumoniae*.

The potential to compare the selected pipelines and their ability to carry out AMR gene detection and prediction was greatly hindered by the lack of standardisation in the output format. This necessitated manual curation of the genes and mutations corresponding to resistance to individual antibiotics, creating an avenue for error. As demonstrated, the AMR genes listed by CARD ontology terms appear difficult to match against annotation pipelines and lead to underperformance in phenotype prediction. The recently developed pipeline hARMonisation has aimed to address this challenge by harmonising the gene names and formats across annotation tools (Mendes et al., 2024). Its application spans widely across 17 bacteria-agnostic annotation tools but is not adapted to report the association of each gene with resistance phenotype beyond what is reported by individual tools. Hence, a gap remains for the creation of a community standardisation tool for gene annotations and AMR association with individual antibiotics. This will be necessary to enable the true application of minimal models.

Importantly, the benchmarking of tools and the demonstration of performance across pipelines and drugs point to clear prediction shortfalls, where known mechanisms do not account for all resistance. These niches present opportunities for the discovery of new mechanisms driving resistance in *K. pneumoniae*. In this dataset, we demonstrate consistent underperformance in predictions of resistance to amikacin, ampicillin/sulbactam, cefepime, cefoxitin, cefuroxime, piperacillin/tazobactam and tetracycline. Interestingly, these findings were not saturated in a single antibiotic class, which can be attributed to several factors. Firstly, although antibiotics belonging to the same class are similar in function, they nevertheless exhibit differences in chemical structures and belong to different development generations. Hence, an antibiotic-to-class approach to prediction might not be sufficient to explain the subtleties of differential resistance to antibiotics in the same class. This was previously commented on by multiple studies which attempted to carry out phenotype prediction (Mahfouz et al. 2020). Hence, the use of gene-to-antibiotic annotations might be more appropriate for the use of minimal models for phenotype prediction. Nevertheless, the same gaps were also identified by tools which carry out gene-to-antibiotic predictions, which supports the value of our model benchmarks.

In addition, the analysis is challenged by some clear limitations of the annotation tools and the fragmented genomes, which resulted in the inability to incorporate the effect of ARG copy into the analysis. Samples in the BV-BRC database were sequenced predominantly using short-read sequencing techniques, resulting in multiple fragmented genomes of many contigs and few fully resolved genomes from long-read or hybrid sequencing. As a result, tandem repeats, which may result in gene amplifications, can be collapsed in the process of genome assembly. Tandem gene amplifications are a known mechanism of modulating resistance in gramnegative bacteria, which was impossible to account for in this analysis (Saathoff et al. 2023). ARG copy number is also dictated by the carriage of plasmids, which is often impossible to resolve. Dosage effects have been hypothesised to have a major role in resistance to carbapenems (Chen et al. 2022).

This analysis is also inevitably biassed by the characteristics of the available dataset. We opted to use the *K. pneumoniae* samples available through the BV-BRS database as it was utilised by multiple previous studies and appears to be emerging as a somewhat standardised dataset of bacterial samples, suitable for model benchmarking (Biffignandi et al. 2024; Zhou et al. 2024). The database presents a large and diverse collection of *K. pneumoniae* samples spanning geographies and sampling niches. However, the collection is often biased in favour of one of the resistance phenotypes, as demonstrated in Figure 1. This poses a challenge for the learning of ML models and could render the ARG features irrelevant. Of the antibiotics we studied, cefuroxime appeared most imbalanced, followed by amikacin; however, cefepime, cefoxitin, tetracycline and piperacillin/tazobactam appear balanced when samples labelled intermediate are converted to resistant. This binarisation of the target label could be a further confounding factor due to changes in MIC breakpoints through time, also noted by Mahfouz et al. (2020). For example, ciprofloxacin CLSI breakpoints for *Enterobacteriaceae* have increased from the initial S ≤ 0.25, I = 0.5, R ≥ 1 to (S) ≤1, intermediate (I) = 2, resistant (R) ≥4. Hence, reference to the source studies and manual curation could be beneficial to ensure consistency in MIC values through the re-assignment of binary labels. This was out of the scope of this study as we aimed to utilise the sample cohort as a standardised dataset. Furthermore, the BV-BRS dataset is also prone to label errors related to testing and reporting practices in different geographies. Hence, further analysis is necessary to estimate the signal-to-noise ratio of resistance for samples in these subsets to truly pinpoint whether the source of underperformance is insufficient knowledge of the biological mechanisms underlying the observed phenotypes.

Once validated as true discovery opportunities, the search for new antimicrobial mechanisms for these antibiotics should include broad variation across the genome, spanning variants in genes as well as variation outside coding regions. Analysis by Mahfouz et al. (2020) discussed the role of regulatory regions in determining the roles of multi-subunit efflux pumps and porins in determining phenotype, showing that the removal of multi-drug-associated genes can increase prediction accuracy. As we did not aim to maximise performance but instead point to where knowledge is missing, we did not carry out any further feature engineering or selection beyond subsetting minimal ARG sets. Therefore, we argue that regulatory sites should be characterised as a class of contributors, rather than excluding the mechanisms they influence. Furthermore, as false positive results are also of major clinical concern, including markers of susceptibility in ARG databases might also be beneficial. In this analysis, we relied on the penalties of the Elastic Net and XGBoost models to carry out appropriate feature selection in the context of the prediction task. Their ability to do so is discussed in depth in the following section, where we also review the function of each antimicrobial, its respective cellular targets and the known resistance mechanisms in *K. pneumoniae*.

### Known resistance mechanisms across antibiotic classes, and opportunities for discovery

#### Aminoglycosides

The aminoglycoside class of antibiotics includes gentamicin, tobramycin and amikacin which were represented in our dataset (Krause et al. 2016). The antibiotics first bind to compounds on the surface of the bacteria - lipopolysaccharide, phospholipids, and outer membrane proteins and induce leakiness. Upon entry into the cell, aminoglycosides affect the function of ribosomal subunits, which causes mistranslation. At sub-inhibitory concentrations, aminoglycosides could also inhibit transcription (Goh et al., 2002). Amikacin specifically has been shown to also lead to inhibition of cell division (Possoz et al., 2007).

Aminoglycoside modifying enzymes, which catalyse inactivation by acetylation (aminoglycoside acetyltransferases, AAC), phosphorylation (aminoglycoside phosphotransferases, APH), or adenylation (aminoglycoside nucleotidyl-transferases, ANT) of the molecule, are the leading cause of the rapid increase and dissemination of resistance across species. The plasmid-carried acetyltransferase *aac(6′)-Ib* or *aacA4* and aminoglycoside adenylyltransferase *aph(3’)-I* confer high resistance to amikacin and other aminoglycosides, but not gentamicin (Chiem et al., 2015; Li et al., 2015). The *aph(3’)-I* genes were present in minimal subsets for all aminoglycosides, even in the cases of ResFinder and RGI, however, the XGBoost model correctly underweighted them in predictions of gentamicin and overweighted them for amikacin resistance. Other 16S-RMTases involved in aminoglycoside resistance are *rmtA*, which in this analysis was given high importance for amikacin classification, r*mtB*, and r*mtC* (Lee et al. 2018). Non-linear resistance to aminoglycosides has been associated with upregulation of the chromosomal *eis* (enhanced intracellular survival) gene caused by mutations in its promoter, which was not annotated in this analysis (Chen et al., 2012).

#### Penicillins

Penicillins are broad-spectrum antibiotics which inhibit the cross-linking of peptidoglycan within the bacterial cell wall (Wilkowske, 1977). The enzymes responsible for the cross-linking of glycan chains are penicillin-binding proteins (PBPs) situated in the periplasm, which are the target for penicillin and other β-lactam antibiotics. As such, mutations in *pbp* genes can lead to resistance to penicillin and other β-lactams, when they lead to a reduction in binding activity (Haenni and Moreillon, 2006). Furthermore, TEM, SHV, CTX-M, VEB, and GES genes encode β-lactamases which hydrolyse the penicillins and other antibiotics of this class by breaking down the β-lactam ring (Canton et al., 2012). *K. pneumoniae* is intrinsically resistant to ampicillin and penicillin due to the chromosomal presence of SHV-1. In line with this, all samples phenotyped for ampicillin were resistant, hence, they were not included in ML predictions (Fig 1).

#### Cephalosporins

The cephalosporin class was represented by most antibiotics in out dataset, encompassing cefazolin, cefepime, cefoxitin, ceftazidime and cefuroxime, which fall into different generations (Xie et al. 2018). Prediction of resistance to these antibiotics differed in performance, which can be attributed in some cases to differences in mechanisms as well as the differences in sample size and class imbalances. Similarly to penicillin, cephalosporins also target PBPs to inhibit peptidoglycan crosslinking. Cephalosporins use porins to cross the outer membrane of gram-negative bacteria. Resistance is mediated by modifications of PBPs, porins, efflux pumps and cephalosporinases (Djoric et al., 2020). The presence of cephalosporinase *acc-1*, a variant of AmpC, was identified in this dataset. Regulatory cascades of AmpC are also associated with different levels or resistance in a non-linear manner (Araten et al. 2023). Extended-spectrum β-lactamases dominated the dataset, such as CTX-M, CMY, DHA, FOX, OXA, TEM, GES.

#### Carbapenems

Carbapenems belong to the broader class of β-lactam antibiotics. They play a critically important role in our antibiotic armamentarium, as they are usually reserved as last resort antibiotics. The antibiotics first enter the cells though outer membrane proteins (OMPs) – porins. Similarly to penicillin, imipenem and other carbapenems interact with PBPs, but unlike penicillin, have some stability against β-lactamases but not dehydropeptidase I (DHP-I). Hence, other carbapenems which are more stable in the presence of dehydropeptidase I were developed, such as meropenem and ertapenem. Resistance is also mediated by preventing peptidoglycan formation through PBP interactions (Montaner et al., 2022). A detailed review of carbapenems is available by Papp-Wallace et al. (2011).

Mechanisms of resistance to carbapenems include the production of β-lactamases, efflux pumps, and mutations that alter the expression and/or function of porins and PBPs (Meletis, 2016). In this dataset, we observed the presence of β-lactamase genes *blaOXA, blaKPC, blaNDM, blaIPM* and *blaVIM*. In addition, the *ompK* gene encoding an outer membrane porin protein was also annotated as a contributor to carbapenem resistance (Kaczmarek et al., 2006). It has been demonstrated that intact OmpK porins are major contributors to susceptibility as they enable efficient diffusion of β-lactams, which cannot be compensated for by β-lactamases, while mutational loss of functional porins confers high levels of resistance (Sugawara et al., 2016). Gene presence is believed to contribute to resistance predominantly linearly and gene copy number has been shown to lead to higher resistance (Abe et al. 2021), which is not currently accounted for in our minimal model.

#### Fluoroquinolones

The dataset encompassed the second-generation fluoroquinolone ciprofloxacin and third-generation levofloxacin (Jacoby and Hooper, 2016). DNA gyrase and topoisomerase IV are the main targets of this antibiotic class, which can differ between species, hence fluoroquinolone action is limited (Correia et al. 2017). This can present a limitation for the majority of annotation tools, as they are species-agnostic. Fluoroquinolones bind and stabilise the complex formed between the cleaved DNA and the type II topoisomerase enzyme. This binding prevents the disassociation of the enzyme-DNA cleavage complex and blocks the ligation process, resulting in the accumulation of these complexes in the cytoplasm. Furthermore, interaction with GyrA and ParC causes enzymes to stall on the DNA and lose their ability to induce strand relaxation or negative supercoiling. Mutations in *gyrA* and *parC*, which render the antibiotic target ineffective, were scored in this dataset. . *Qnr* variants which play a further role in resistance by protecting topoisomerases, were also abundant in the dataset (Jacoby et al. 2015). They fall within the broader spectrum of plasmid-mediated quinolone resistance (PMQR) genes. Finally, modifications in efflux pumps and porins also contribute to resistance by modulating cellular entry. For example, the OqxAB efflux pump, when associated with mobile genetic elements, appears overexpressed and confers resistance to quinolones (Wyres et al., 2020). This mechanism presents an example of a locus acquired by *K. pneumoniae* and disseminated to other species (Li et al. 2019).

#### Combination therapies

Three combination therapies were represented in this dataset - trimethoprim/sulfamethoxazole, ampicillin/sulbactam and piperacillin/tazobactam. The strategy behind using combination therapies as antimicrobials is to target multiple independent systems at once, which should challenge the development of resistance. Hence, the presence of multiple resistance mechanisms is needed to confer resistance to a combination therapy, causing non-linearity of the inputs and the labels in our models. For example, the antibiotic trimethoprim belongs to the diaminopyrimidines class, while sulfamethoxazole represents sulphonamides. In this case, diaminopyrimidines target cofactor biosynthesis by inhibiting dihydrofolate reductase (DHFR) enzymes. Resistance mechanisms are associated with accumulation of mutations in *dfrC* genes, encoding these enzymes. Sulfonamides also inhibit cofactor biosynthesis but through the alternative mechanism of competitively replacing *p*-aminobenzoic acid in the synthesis of folic acid. Resistance to this class is mediated by mutations in *folP* or through the acquisition of *sul* genes, which encode sulfa-insensitive, divergent DHPS enzymes, an alternative to the usual trimethoprim/sulfonamide target (Vankatesan et al, 2023). High resistance accuracy in this dataset was achieved by annotating the three genes *sul1, sul2* and *sul3*. However, this is likely due to the complete correlation of resistance to trimethoprim and sulfamethoxazole in this dataset, also linked to the historic use of trimethoprim solely in combination with sulfamethoxazole (Hamilton-Miller, 1979).

## Conclusions

In this analysis, we utilised minimal models of antibiotic resistance in *K. pneumoniae* to identify the degree to which known mechanisms account for phenotypic resistance to the antibiotics amikacin, ampicillin/sulbactam, cefepime, cefoxitin, cefuroxime, piperacillin/tazobactam and tetracycline. This allowed us to determine the opportunity for discovery of new AMR variants and mechanisms. We also show that resistances to gentamicin, imipenem and ertapenem are well-classified based on the reviewed annotations, hence, new ML models can be benchmarked against this dataset and are expected to find well known AMR markers. We utilised eight commonly used AMR annotation tools to identify known mechanisms of resistance in the sample cohort and demonstrated stark differences in the breadth of annotations and prediction potential, ultimately showing that no tool is consistently better than any other. We note that the use of minimal models would ultimately be unlocked when minimal gene sets are annotated in a gene-to-antibiotic manner, as opposed to the present gene-to-class annotation provided by most tools, to capture subtleties of differential resistance to antibiotics in the same class. We discussed the major known AMR genes and mechanisms in this dataset with respect to the bacterium *K. pneumoniae*. Further work will seek to analyse the noise-to-signal ratio in this dataset and validate the discovery opportunities for further whole genome ML-enabled mining. Ultimately, the analysis presents a valuable exploration of the BV-BCR dataset as a standardised dataset for benchmarking and developing novel ML models, and niches where they can be validated or used to discover unknown markers.

## Author Declarations

### Data availability

The datasets generated and/or analysed during the current study are available in the ‘BV-BRC Klebsiella pneumoniae AMR Annotations’ repository, https://zenodo.org/records/14126594?token=eyJhbGciOiJIUzUxMiJ9.eyJpZCI6IjQwZDI1Yz U2LTU1NzYtNDdiNy1iYzdmLWRjNzQ1YmE1NDljNyIsImRhdGEiOnt9LCJyYW5kb20i OiI4NDZmMzgyM2E5OGEwMWY0ODY0MmFkOTU1YWU2OTI4YiJ9.jreefdp3-X4bLG2cI2lWPm7tkpzBWN05fEY1Ol-9Ht54kCJECsCwAt96zx8p7wzUR3hURMU9KduJ7uizUCmr1A

### Funding

The research was conducted as a part of the PhD studentship funded by The Ineos Oxford Institute for antimicrobial research (IOI)

### Authors’ contributions

K.K. wrote the main manuscript, conducted bioinformatic analysis of the data and prepared the figures; C.C. and J.P. reviewed the manuscript and advised the analysis

## Acknowledgements

We would like to acknowledge the contributions of Sebastian Bruchmann who provided invaluable feedback on the final manuscript.

## Notes

https://zenodo.org/records/14126594?token=eyJhbGciOiJIUzUxMiJ9.eyJpZCI6IjQwZDI1YzU2LTU1NzYtNDdiNy1iYzdmLWRjNzQ1YmE1NDljNyIsImRhdGEiOnt9LCJyYW5kb20iOiI4NDZmMzgyM2E5OGEwMWY0ODY0MmFkOTU1YWU2OTI4YiJ9.jreefdp3-X4bLG2cI2lWPm7tkpzBWN05fEY1Ol-9Ht54kCJECsCwAt96zx8p7wzUR3hURMU9KduJ7uizUCmr1A

### Competing Interest Statement

The authors have declared no competing interest.

### Summary of Updates

This revision has only changed the author's order.

