## Supplementary Materials for "Finding the known unknowns: minimal machine learning models of resistance identify novel antibiotic resistance discovery opportunities in *Klebsiella pneumoniae*"

| Antibiotic | Antibiotic class | Abbreviation |
|---|---|---|
| amikacin | aminoglicosides | AMK |
| ampicillin | penicillins | AMP |
| aztreonam | monobactams | ATM |
| ampicillin/sulbactam | penicillins and beta-lactams | SAM |
| cefazolin | cephalosporins | CFZ |
| cefepime | cephalosporins | FEP |
| cefoxitin | cephalosporins | FOX |
| ceftazidime | cephalosporins | CAZ |
| ceftriaxone | cephalosporins | CRO |
| cefuroxime | cephalosporins | CXM |
| ciprofloxacin | fluoroquinolones | CIP |
| ertapenem | carbapenems | ETP |
| gentamicin | aminoglicosides | GEN |
| imipenem | carbapenems | IPM |
| levofloxacin | fluoroquinolones | LVX |
| meropenem | carbapenems | MEM |
| piperacillin/tazobactam | penicillins and beta-lactams | TZP |
| tetracycline | tetracyclines | TET |
| tobramycin | aminoglicosides | TOB |
| trimethoprim/sulfamethoxazole | sulfonamides and diaminopyrimidines | SXT |

*Supplementary Table 1: List of antibiotics and their classes, used to match them against annotation tools, and their abbreviations.*

| Tool | Organism | Input | Database | Output |
|---|---|---|---|---|
| **ABRicate** | Agnostic | Contigs | NCBI, CARD, ARG-ANNOT, Resfinder, MEGARES, EcOH, PlasmidFinder, Ecoli_VF and VFDB | gene-to-class |
| **AMRFinderPlus** | Agnostic | Protein and/or assembled nucleotide sequences | AMRFinderPlus | gene-to-class |
| **DeepARG** | Agnostic | Metagenomic data, Short reads, Lond sequences | DeepARG-DB | gene-to-class |
| **ResFinder** | Agnostic | Contigs | ResFInder, PointFinder, DesintFinder | gene-to-antibiotic |
| **RGI** | Agnostic | Contigs | CARD | gene-to-antibiotic |
| **SraX** | Agnostic | Contigs | AMR DB | gene-to-class |
| **StarAMR** | Agnostic | Contigs | ResFinder, PointFinder, and PlasmidFinder | gene-to-antibiotic |
| **Kleborate** | *Klebsiella pneumoniae* | Contigs | Kleborate | gene-to-class |
| **AMRplusPlus** | Agnostic | Metagenomic data | MEGARes | not reported |
| **Ariba** | Agnostic | Genome reads | ARG-ANNOT, CARD, MEGAres and ResFinder | gene-to-class |
| **C-sstar** | Agnostic | Contigs | ARG-ANNOT, ResFinder or ResGANNOT | gene names with no association |
| **Groot** | Agnostic | Metagenomic data | CARD, Resfinder, ARG-ANNOT | not reported |
| **KmerResistance** | Agnostic | Contaminated and poorly sequenced samples | Any indexed with KMA | not reported |
| **Mykrobe** | *Mycobacterium tuberculosis, Staphylococcus aureus, Shigella sonnei, Salmonella typhi and Salmonella enterica serotype Paratyphi B.* | Genome reads | Mykrobe | gene-to-antibiotic |
| **PointFinder** | *Salmonella enterica, Escherichia coli and Campylobacter jejuni* | Contigs | PointFinder | gene-to-class |
| **ResFams** | Agnostic | Contigs | Resfams | gene-to-antibiotic |
| **SRTS2** | Agnostic | Genome reads | SRST2 | gene-to-class |
| **TBProfiler** | *Mycobacterium tuberculosis* | Genome reads | tbdb database | gene-to-antibiotic |
| **fARGene** | Agnostic | Fragmented metagenomic data or longer sequences | fARGene | gene-to-class |

*Supplementary table 2: List of the 19 AMR gene annotation tools identified for this analysis and description of their input, default databases and output formats. The eight tools which were compared in this analysis are listed above the bold divider.*

A)

| Tool | AMK | AMP | SAM | ATM | CFZ | FEP | FOX | CAZ | CRO | CXM | CIP | ETP | GEN | IPM | LVX | MEM | TZP | TET | TOB | SXT |
|---|---|---|---|---|---|---|---|---|---|---|---|---|---|---|---|---|---|---|---|---|
| RGI XGBoost | 0.971 | 0 | 0.083 | 0 | 0 | 0 | 0.001 | 0 | 0 | 0 | 0.775 | 0.004 | 0.961 | 0.992 | 0.852 | 0.983 | 0.029 | 0.968 | 0.723 | 0.779 |
| RGI linear | 0.969 | 0 | 0.005 | 0 | 0 | 0.003 | 0.001 | 0 | 0.006 | 0 | 0.761 | 0.004 | 0.93 | 0.992 | 0.857 | 0.983 | 0.018 | 0.97 | 0.723 | 0.822 |
| ResFinder XGBoost | 0.911 | 0 | 0.512 | 0.655 | 0 | 0.558 | 0.953 | 0.878 | 0.372 | 0 | 0 | 0.992 | 0.961 | 0.943 | 0 | 0.928 | 0.522 | 0.943 | 0.723 | 0.766 |
| ResFinder linear | 0.904 | 0 | 0.302 | 0.127 | 0 | 0.544 | 0.953 | 0.348 | 0.362 | 0 | 0.002 | 0.921 | 0.931 | 0.943 | 0 | 0.914 | 0.587 | 0.942 | 0.721 | 0.817 |
| StarAMR XGBoost | 0.904 | 0 | 0.454 | 0.659 | 0 | 0.551 | 0.953 | 0.859 | 0.113 | 0 | 0.002 | 0.992 | 0.961 | 0.944 | 0 | 0.915 | 0.523 | 0.943 | 0.721 | 0.769 |
| StarAMR linear | 0.899 | 0 | 0.332 | 0.638 | 0 | 0.55 | 0.953 | 0.829 | 0.125 | 0 | 0.005 | 0.992 | 0.931 | 0.944 | 0 | 0.914 | 0.54 | 0.943 | 0.721 | 0.817 |
| AMRFinderPlus XGBoost | 0.912 | 0 | 0.04 | 0.021 | 0.599 | 0.517 | 0.458 | 0.423 | 0.474 | 0.008 | 0.785 | 0.979 | 0.966 | 0.944 | 0.916 | 0.918 | 0.137 | 0.943 | 0.905 | 0.83 |
| AMRFinderPlus linear | 0.912 | 0 | 0.074 | 0.041 | 0.559 | 0.389 | 0.481 | 0.469 | 0.544 | 0.058 | 0.771 | 0.979 | 0.963 | 0.944 | 0.91 | 0.912 | 0.173 | 0.943 | 0.905 | 0.83 |
| DeepARG XGBoost | 0.909 | 0 | 0.517 | 0.282 | 0.701 | 0.458 | 0.698 | 0.535 | 0.681 | 0.083 | 0.286 | 0.312 | 0.93 | 0.794 | 0.336 | 0.905 | 0.479 | 0.803 | 0.669 | 0.138 |
| DeepARG linear | 0.9 | 0 | 0.42 | 0.186 | 0.631 | 0.431 | 0.669 | 0.479 | 0.637 | 0.025 | 0.24 | 0.206 | 0.932 | 0.813 | 0.226 | 0.923 | 0.464 | 0.833 | 0.576 | 0.111 |
| Abricate XGBoost | 0.922 | 0 | 0.049 | 0 | 0.885 | 0.562 | 0.758 | 0.834 | 0.847 | 0.008 | 0 | 0.909 | 0.953 | 0.949 | 0.064 | 0.946 | 0.581 | 0.962 | 0.736 | 0.833 |
| Abricate linear | 0.922 | 0 | 0.127 | 0.007 | 0.773 | 0.553 | 0.764 | 0.71 | 0.769 | 0.075 | 0.003 | 0.854 | 0.951 | 0.941 | 0.051 | 0.939 | 0.581 | 0.959 | 0.706 | 0.833 |
| SraX XGBoost | 0.929 | 0 | 0 | 0 | 0 | 0.6 | 0.332 | 0.012 | 0 | 0 | 0.115 | 0.277 | 0.963 | 0.762 | 0.253 | 0.794 | 0 | 0.452 | 0.643 | 0.683 |
| SraX linear | 0.922 | 0 | 0 | 0 | 0 | 0.45 | 0.334 | 0.002 | 0 | 0 | 0.055 | 0.438 | 0.957 | 0.756 | 0.094 | 0.747 | 0 | 0.451 | 0.649 | 0.343 |
| Kleborate XGboost | 0.926 | 0 | 0.473 | 0 | 0.888 | 0.55 | 0.779 | 0.827 | 0.894 | 0.05 | 0.797 | 0.963 | 0.955 | 0.944 | 0 | 0.917 | 0.586 | 0.942 | 0.806 | 0.789 |
| Kleborate linear | 0.916 | 0 | 0.366 | 0 | 0.71 | 0.572 | 0.765 | 0.662 | 0.738 | 0.033 | 0.782 | 0.933 | 0.947 | 0.928 | 0 | 0.894 | 0.597 | 0.906 | 0.808 | 0.777 |

B)

| Tool | AMK | AMP | SAM | ATM | CFZ | FEP | FOX | CAZ | CRO | CXM | CIP | ETP | GEN | IPM | LVX | MEM | TZP | TET | TOB | SXT |
|---|---|---|---|---|---|---|---|---|---|---|---|---|---|---|---|---|---|---|---|---|
| RGI XGBoost | 0.271 | 1 | 0.98 | 1 | 1 | 1 | 0.997 | 0.998 | 1 | 1 | 0.96 | 0.993 | 0.846 | 0.056 | 0.853 | 0.171 | 0.992 | 0.584 | 0.885 | 0.955 |
| RGI linear | 0.289 | 1 | 0.998 | 1 | 1 | 1 | 0.997 | 0.998 | 1 | 1 | 0.962 | 0.994 | 0.856 | 0.056 | 0.856 | 0.171 | 0.993 | 0.587 | 0.89 | 0.943 |
| ResFinder XGBoost | 0.788 | 1 | 0.96 | 0.935 | 0 | 0.927 | 0.609 | 0.931 | 0.982 | 0 | 1 | 0.767 | 0.853 | 0.879 | 1 | 0.867 | 0.925 | 0.589 | 0.894 | 0.959 |
| ResFinder linear | 0.793 | 1 | 0.974 | 0.995 | 0 | 0.92 | 0.615 | 0.979 | 0.98 | 0 | 0.998 | 0.786 | 0.861 | 0.873 | 1 | 0.879 | 0.872 | 0.592 | 0.898 | 0.941 |
| StarAMR XGBoost | 0.743 | 1 | 0.958 | 0.922 | 0 | 0.918 | 0.601 | 0.921 | 0.992 | 0 | 0.999 | 0.745 | 0.851 | 0.843 | 0 | 0.868 | 0.911 | 0.588 | 0.9 | 0.958 |
| StarAMR linear | 0.761 | 1 | 0.983 | 0.936 | 0 | 0.92 | 0.612 | 0.932 | 0.993 | 0 | 0.998 | 0.752 | 0.859 | 0.883 | 0 | 0.874 | 0.897 | 0.591 | 0.902 | 0.941 |
| AMRFinderPlus XGBoost | 0.639 | 1 | 0.991 | 0.976 | 0.925 | 0.883 | 0.788 | 0.961 | 0.946 | 0.998 | 0.954 | 0.91 | 0.075 | 0.847 | 0.925 | 0.861 | 0.946 | 0.591 | 0.752 | 0.886 |
| AMRFinderPlus linear | 0.643 | 1 | 0.987 | 0.965 | 0.946 | 0.898 | 0.795 | 0.964 | 0.956 | 0.996 | 0.958 | 0.921 | 0.082 | 0.873 | 0.931 | 0.875 | 0.943 | 0.594 | 0.756 | 0.886 |
| DeepARG XGBoost | 0.421 | 1 | 0.967 | 0.944 | 0.94 | 0.901 | 0.761 | 0.955 | 0.959 | 0.989 | 0.905 | 0.813 | 0.705 | 0.501 | 0.917 | 0.764 | 0.859 | 0.701 | 0.808 | 0.953 |
| DeepARG linear | 0.468 | 1 | 0.968 | 0.949 | 0.952 | 0.863 | 0.78 | 0.962 | 0.968 | 0.995 | 0.931 | 0.866 | 0.714 | 0.466 | 0.934 | 0.77 | 0.874 | 0.676 | 0.833 | 0.958 |
| Abricate XGBoost | 0.607 | 1 | 0.992 | 1 | 0.928 | 0.91 | 0.758 | 0.931 | 0.924 | 0.997 | 0.995 | 0.751 | 0.866 | 0.85 | 0.97 | 0.808 | 0.893 | 0.614 | 0.879 | 0.874 |
| Abricate linear | 0.705 | 1 | 0.987 | 0.994 | 0.954 | 0.889 | 0.774 | 0.954 | 0.952 | 0.994 | 0.999 | 0.78 | 0.868 | 0.861 | 0.989 | 0.826 | 0.898 | 0.6 | 0.91 | 0.874 |
| SraX XGBoost | 0.651 | 1 | 1 | 0 | 1 | 0.803 | 0.868 | 0.992 | 1 | 1 | 0.968 | 0.762 | 0.868 | 0.342 | 0.941 | 0.398 | 0 | 0.768 | 0.859 | 0.789 |
| SraX linear | 0.667 | 1 | 1 | 0 | 1 | 0.828 | 0.86 | 0.996 | 0.998 | 1 | 0.98 | 0.769 | 0.874 | 0.351 | 0.969 | 0.438 | 0 | 0.77 | 0.843 | 0.886 |
| Kleborate XGboost | 0.635 | 1 | 0.966 | 0 | 0.923 | 0.906 | 0.749 | 0.94 | 0.926 | 0.991 | 0.95 | 0.918 | 0.838 | 0.845 | 0 | 0.862 | 0.875 | 0.623 | 0.883 | 0.951 |
| Kleborate linear | 0.661 | 1 | 0.973 | 0 | 0.956 | 0.859 | 0.779 | 0.958 | 0.958 | 0.995 | 0.947 | 0.916 | 0.842 | 0.868 | 0 | 0.851 | 0.866 | 0.66 | 0.902 | 0.954 |

*Supplementary Table 3: A) Specificity and B) Sensitivity of phenotype predictions across antibiotics and annotation pipelines. Gene-to-antibiotic predictions are placed above the bold table line and gene-to-class predictions are placed below. Ampicillin (AMP) performance sensitivity and specificity are determined by all samples being resistant fort this antibiotic.*
